# Spatiotemporal mapping of the neural markers of prediction error processing across multisensory and unisensory modalities

**DOI:** 10.1101/2022.02.11.480053

**Authors:** Priyanka Ghosh, Siddharth Talwar, Arpan Banerjee

**Affiliations:** Cognitive Brain Dynamics Lab, National Brain Research Centre, NH-8, Manesar, Gurgaon, Haryana-122052, India

**Author notes:** Authors having equal contributions, co-first authors (Priyanka Ghosh, Siddharth Talwar). Author’s present address: Crossmodal Perception and Plasticity (CPP) Lab, Université Catholique de Louvain (UCL), Place Cardinal Mercier 10, 1348 Louvain-la-Neuve, Belgium.

**Keywords:** Prediction error, MMN, P300, Multisensory facilitation, Source Localization

## Abstract

Prediction errors in the brain are indexed by two event-related potentials – MMN and P300, which are elicited upon violation of regularity in the occurrence of repetitive stimuli. While MMN reflects the brain’s ability to perform automatic comparisons between consecutive stimuli and provides an electrophysiological index of sensory error detection, P300 is associated with cognitive processes such as update in working memory. Till date, there has been extensive research on the roles of MMN and P300 individually, because of their potential to be used as clinical markers of consciousness and attention, respectively. However, the relationship between these two ERPs, specifically in terms of their underlying cortical generators, in context of prediction error propagation along the hierarchical brain across multiple modalities is an open question. Our objective in this article is two-fold. First, we reconfirm previous reports regarding the generators of MMN and P300 in sensor space through source-space analysis using an accurate individual subject level co-registration of MRI and EEG data collected from healthy humans. We demonstrate that in multisensory environments, MMN and P300 markers represent “modality-specific” and “modality-independent” information processing, respectively. Advancing an earlier understanding that multisensory contexts speed up *early sensory processing*, our study reveals that this temporal facilitation extends to even the later components of *prediction error processing*, using custom-designed experiments that allow comparisons across different modality combinations. Such knowledge can be of immense value in clinical research for determining the stages of various treatments in aging, schizophrenia and depression, and their efficacy on cognitive function.

## INTRODUCTION

The brain attends to complex stimuli by minimizing its response to frequent events that do not require extensive processing thus, freeing up the cognitive resources to process unexpected events (Fontolan et al., 2014; Friston, 2005; Aitchison & Lengyel, 2017). To subserve this goal, a neural representation (internal model) is constructed based on prior information from previous sensory inputs and any mismatch between the model and the subsequent sensory input gives rise to a “prediction error” (Kok et al., Winkler and Czigler, 2012). The existence of prediction error representation at the observational scale can be taken as the evidence of a computational/algorithmic framework of predictive coding principles (Aitchison & Lengyel, 2017), although by itself that doesn’t divulge the specific nature of the internal model. At the level of ERPs in the brain, prediction error can be captured as the MMN (Mismatch Negativity) and the P300 component (Aitchison & Lengyel, 2017; Banellis et al., 2020; Calcus et al., 2015; Chennu et al., 2013; Stefanics et al., 2014). Thus, the study of these two prediction error signals under multisensory contexts can aid in our understanding of how the brain scans the environment to distinguish deviation from regularity, specifically the temporal stages of the prediction process and their underlying cortical sources. Prediction error propagation is organized hierarchically, such that the prediction error arising from a given area in turn serves as the input to the next area (Friston, 2005), consequently updating the internal models at the hierarchical stages (Wacongne et al., 2011). Hence, to study the spatiotemporal representation of prediction error processing across sensory modalities, we use an oddball paradigm, the most widely used paradigm because of its simplicity, reproducibility, and applicability, where repetitive ‘standard’ stimuli are interspersed with rare ‘deviant’ stimuli.

### A brief history of MMN and P300

The foundation of MMN and P300 research was laid more than half a century ago where the two ERPs were recognized as the markers of “uncertainty” (Sutton et al., 1965). Since then, both these markers have been extensively studied under various stimulus conditions and clinical scenarios.

MMN appears as a negative deflection in the difference waveform between deviant and standard EEG signals around 100-250 ms post onset of a deviant stimulus with frontal/central/temporal generators in the brain (Garrido et al., 2009; Näätänen et al., 1978, 2007; Näätänen & Michie, 1979; Sams et al., 1985). It is known to index any deviance from a recognized pattern in sensory stimuli, with its amplitude indicating the degree of salience of the prediction error (Picton et al., 2000; Winkler, 2008). P300, on the other hand, is a positive deflection in voltage with a latency between 250 to 500 ms distributed across the fronto-central scalp locations indexing the possible attention switch and conscious perception of stimulus change (Giard et al., 1990; Polich, 2007a; Sutton et al., 1967). Its amplitude is sensitive to oddball probability and the amount of deviance from the standard stimulus, and its latency reflects oddball evaluation time (Johnson & Donchin, 1980). The MMN belongs to a group of ERPs referred to as N200 and can be divided into N2a or MMN, N2b and N2c (Patel and Azzam 2005; Folstein and Van Petten, 2008). N2a can be elicited by both attended and unattended stimuli, whereas the N2b and N2c only occur when attention is directed to target stimuli (Folstein & Van Petten, 2008; Muller-Gass et al., 2005). The P300 wave too, is further separable into the slightly earlier (250-300ms), fronto-central P3a and the later (300-350ms), centro-parietal P3b, thought to be evoked by unpredictable/ task-irrelevant distractors and rare-target/ task-relevant stimuli, respectively (Courchesne et al., 1975; A. Kok, 2001; Polich & Criado, 2006; Squires et al., 1975). Our study, however, includes the temporal and spatial properties of only P3b as we use a two stimuli oddball paradigm where the oddball is the target. P300 being a late component, is associated with updating the working memory in oddball counting tasks (Donchin, 1981; Donchin & Coles, 1988) and has been linked to decision making processes as well (Rohrbaugh et al., 1974). P3b is particularly thought to be involved in the deployment of selective attention to task-relevant stimuli under conscious awareness (Kok, 2001; Polich and Criado, 2006). On the contrary, MMN can be elicited pre-attentively, during non-attentive states such as sleep (Molholm et al., 2005), behavioral unconsciousness (Atienza et al., 2001; Koelsch et al., 2006), or even in coma (Morlet & Fischer, 2014). It has provided researchers access to pre-conscious processing of temporal structure in mostly auditory information beyond the basic sensory stage before it enters conscious perception or stages of attention (Chennu & Bekinschtein, 2012; Wijnen et al., 2007). It however, can also be modulated by attention (Alain & Woods, 1997; Woldorff et al., 1998), the reason why there is a sustained empirical interest in MMN till date. Earlier, MMN was thought to reflect auditory mechanisms only (Nyman et al., 1990) but substantial pieces of evidence now suggest it has a visual counterpart too, popularly known as vMMN (Pazo-Alvarez et al., 2003; Stefanics et al., 2014).

### Importance of research on multisensory prediction errors

Most prediction mismatches in our surroundings are perceived through not one but multiple sensory modalities (Luca et al., 2009), e.g. noticing the brake lights of a car in front while engaged in listening to the radio. From a predictive coding perspective, the internal mental model of sight, sound, smell, taste and touch are integrated with our existing cognitive schemata (Talsma, 2015). Processing of novel bottom-up stimuli in one modality can subsequently modify the neural representation and hence top-down prediction, of a stimulus in another sensory modality. If multisensory integration involves such complex interactions between top-down and bottom-up processes, then it should take place at multiple stages of processing, based on the complexity of the stimuli involved (Molholm et al., 2002). The first distinct stage of multisensory integration was identified at about 100 ms of speech stimulus onset where the audio-visual N1 component peaked earlier than that of the auditory speech stimulus alone (Stekelenburg & Vroomen, 2007; Van Wassenhove et al., 2005). Interestingly, a few studies using oddball stimuli have also shown that multimodal stimulation (visual-audio-tactile and visual-audio) induces a significant early onset of latency (Marucci et al., 2021; Stefanics et al., 2005), and a significant increase in the amplitude of the P300 potentials (Fleming et al., 2020; Marucci et al., 2021) as compared to the corresponding unimodal stimulations. Feng et al.(2008), however, did not observe such multisensory benefits for P300 latencies along with Giard and Peronnet (1999) who reported that multimodal P300 may not necessarily be a linear sum of unimodal P300 components. In a recent study using the oddball paradigm (Shiramatsu et al., 2021), the latency of audio-visual MMN resembled closely to the latency of audio MMN than that of visual MMN, indicating the advantage of audio MMN over visual MMN. On the contrary, a previous study (Sittiprapaporn, 2012) using a different oddball paradigm showed visual and audio-visual MMN had shorter latencies than auditory MMN. Thus, the properties of both MMN and P300 based on latency and amplitude vary with the paradigm and experimental parameters like length of the stimulus, stimulus probability, target to target interval and discrimination difficulty (Gonsalvez et al., 2007; Gonsalvez & Polich, 2002; Magliero et al., 1984; Patel & Azzam, 2005; Polich, 2007) as well as physiological variables such as age, attention and neurophysiological disorders (Blackwood, 2000; Dinteren et al., 2014; Erickson et al., 2016; Polich et al., 1990). A critical question that still remains is if the modality of the prediction error changes, whether a common set of neural areas generate these ERPs, or are they an outcome of information processing within the modality-specific sensory regions?

### On what is yet to be known

Despite a huge body of research on MMN and P300, a single comprehensive study comparing the spatiotemporal properties of these two prediction error markers for unisensory and multisensory modalities is not available. From this perspective, we hypothesize that (i) the speeded responses to multisensory stimuli are also seen in the middle/late processing stages of prediction errors, i.e., for MMN/P300, and (ii) there exists a sensory-cognitive dissociation in the source distribution of MMN and P300. To validate our hypotheses, we investigate the properties of prediction error markers - MMN and P300, from very simplistic active viewing and listening tasks using a multimodal oddball paradigm. Furthermore, we perform rigorous source analysis using co-registration with individual subject’s MRI data to reveal the overlapping cortical networks and specialized brain regions specific to unimodal/ multimodal MMN and P300. Traditional oddball paradigms consist of a mismatch between frequent and deviant stimuli belonging to the “same modality”. However, a “cross-modality”, where the modality of the frequent and the deviant stimuli are different, may recruit different brain regions for oddball processing as the non-target standard modality can be completely ignored, unlike the usual multisensory interactions. The goal of the current article is to replicate previous findings of MMN and P300 research in the light of prediction error processing with conceptual advancement in the organization of the cortical sources along the predictive coding hierarchy in the brain.

## METHODS

### Participants

22 healthy volunteers (9 males and 13 females) in the age group of 22 to 43 (mean=25.7, SD=±4.19) years participated in the study. They had a normal or corrected-to-normal vision and were right-handed. All participants had University degrees or higher and reported no history of neurological or audiological problems. They were requested to avoid the intake of any medication/stimulant (e.g., sedative, coffee, etc.) before the experiment. They provided informed written consent at the beginning of the experiment and were remunerated for their participation. The study was carried out following the ethical guidelines and prior approval of the Institutional Human Ethics Committee of the National Brain Research Centre, India.

### Stimuli

The experiment consisted of five different conditions and each condition consisted of two categories of stimuli, i.e., repetitive/frequent/standard and non-repetitive/oddball/deviant. Two of the five conditions presented were ***unimodal***, i.e., audio only and visual only; the third was ***bimodal***, i.e., audio-visual; and the remaining two were ***cross-modal*** in nature. In the first three conditions, the standard and the oddball stimuli were of the same sensory modality which means that the *audio-only condition* comprised of an audio standard and an audio oddball stimuli; the *visual only condition* comprised of a visual standard and a visual oddball stimuli, and; the *audio-visual (AV) condition* comprised of an audio-visual standard and an audio-visual oddball stimuli. The remaining 2 conditions, namely, the *cross-audio* consisted of an audio deviant and a visual standard and the *cross-visual* consisted of a visual oddball and an audio repetitive stimuli. The contents of each condition are tabulated in **Table 1**. Each condition consisted of 400 trials, out of which oddballs constituted 14 percent of the total trials. Each condition was presented in the form of a block of 100 stimuli, where the standard and oddball stimuli of a particular condition were presented in random order and the number of oddball stimuli varied across each block. There were 20 such blocks (5 conditions x 4 blocks) that were randomized and presented to the participants. The participants were prompted about the modality of the upcoming block before every run and to engage their attention throughout the entire block, they were asked to count the number of oddballs presented in each block and report at the end of the block.

**Table 1.** The table lists the standard and the deviant stimuli used in our oddball paradigm for all the five sensory modality conditions.

| **Condition** | **Standard / Frequent / Repetitive** | **Deviant / Oddball / Infrequent / Non-repetitive** |
| --- | --- | --- |
| *Unimodal (Audio only)* | 261.6 Hz | 523.3 Hz |
| *Unimodal (Visual only)* | Blue Square | Red Triangle |
| *Bimodal (Audio-Visual)* | 261.6 Hz and Blue Square | 523.3 Hz and Red Triangle |
| *Cross-modal (Audio deviant)* | Blue Square | 523.3 Hz |
| *Cross-modal (Visual deviant)* | 261.6 Hz | Red Triangle |

The stimuli of the visual only condition consisted of a standard blue square and a deviant red triangle. The auditory stimuli were inspired from musical notes, the standard as the C4 note and the deviant as the C5 note (higher octave), according to the tuning of the A440 pitch standard. All stimuli were presented on a white background on a 21″ LED screen (1280 × 1024 pixels). The participants were asked to keep their eyes open and fixate on a central cross on the screen during the presentation of all auditory stimuli. The inter-stimulus interval also consisted of the same cross-fixation. The length of each oddball and standard stimulus was 350 ms and the inter-trial interval ranged between 800-1200 ms (mean=1000 ms). AV and cross-modal conditions were constructed from combinations of the audio only and the visual only stimuli (listed in Table 1).

### Data Acquisition

EEG was recorded with 64 Ag/AgCl electrodes, using a Neuroscan system (Compumedics NeuroScan, SynAmps2). The electrodes were attached to an elastic cap in a 10−20 International system montage. The data were acquired at a sampling rate of 1000 Hz with the default reference near Cz, grounded to electrode AFz. During the experiment, the participants were seated comfortably at a distance of 60-70cm from the monitor in a sound attenuated and dimly lit room. The participants were requested to make minimal body movements and blink normally during the experiment. The impedance of all electrodes was initially set below 5kΩ and was also monitored in between blocks. Additionally, head digitization was performed using Polhemus Fastrak (Polhemus Inc.) to mark the position of the electrodes and the fiducials based on the placement of the cap on individual participants at the end of the entire EEG session. Individual T1-weighted structural images (MPRAGE) were also obtained using a 3.0 T Philips Achieva MRI scanner (TR = 8.4 ms, FOV = 250 × 230 × 170, flip angle = 8°).

### Preprocessing

EEG was acquired from 22 participants out of which data from 1 participant was discarded due to noisy recordings. The raw data of the remaining 21 participants were imported and each block was epoched. A high pass filter of 0.1 Hz was applied to the data to remove slow drifts in the signal. The data were visually inspected further and 1 channel (F4 in 3 subjects) was interpolated to neighboring electrodes. To identify and remove blink artifacts from the data, independent component analysis (ICA) was employed for each block. Only the blink component obtained as independent component (IC) from eye regions was visually identified and subsequently that IC was rejected. Each block was further epoched to trials of [-500 550] ms where 0 ms marked the onset of the stimulus. The trials were further divided into standard and oddball categories. Subsequently, all trials were subjected to a low pass filter of <45 Hz, baseline correction was applied and the data was re-referenced to linked mastoids. Trials with signal amplitude greater than 100µV and lesser than −100µV were removed and at most 6% of all oddball trials were discarded per subject. To equate the number of standard and oddball trials from each participant, we chose the minimum number of trials that survived artifact rejection in any condition and those many trials were randomly sampled from each condition for each participant. For all conditions, 885 trials each for deviant and standard categories were used. All analyses were done using the MATLAB based FieldTrip toolbox developed at Donders Institute for Brain, Cognition and Behaviour, in Nijmegen, Netherlands (Oostenveld et al., 2011) and custom-written scripts in MATLAB (www.mathworks.com).

### Extracting MMN and P300 peaks

The MMN and P300 peaks were identified for the audio (A), visual (V) and audio-visual (AV) conditions by subtracting the standard waveforms from their respective deviant waveforms. The group-averaged difference waveforms with prominent MMN and P300 peaks and their corresponding topoplots are shown for each condition in **Figure 1**. In an attempt to eliminate researchers’ degrees of freedom and control for Type I errors in our analyses, we used apriori defined temporal windows of interest for MMN/P300 peaks based on previous ERP literature. The largest positive deflection between 250ms to 500ms from the onset of stimulus was considered as a P300 activity and the largest negative deflection between 120ms to 250ms in the difference waveform was considered as an MMN response (Garrido et al., 2009; Näätänen et al., 2007, 1978).

**Figure 1.**
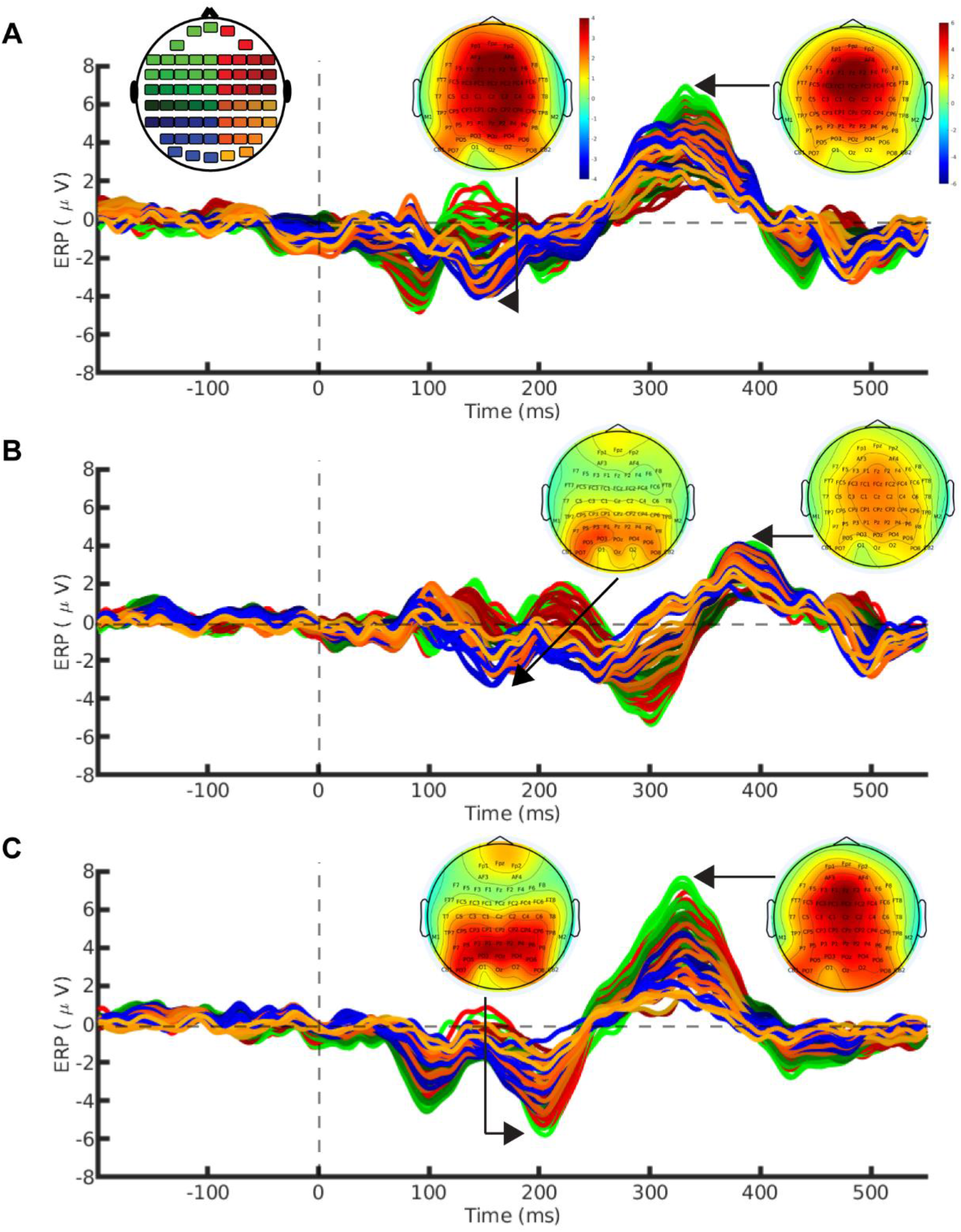
Grand average of difference ERPs (deviant-standard) plotted across 21 subjects for conditions **(A) audio only, (B). visual only** and **(C) audio-visual**. Stimulus onset is at 0ms and the activity before that [-200 ms to 0 ms] corresponds to the pre-stimulus baseline. The topoplots indicate the distribution of the respective MMN and P300 peaks in the sensor-space. The topoplot on the top-left displays the color code assigned to respective scalp channel locations used for plotting the ERPs in A, B and C.

Given that we have 64 sensors to choose from, it is difficult to know the spatial regions of interest (ROI) that would best characterize the MMN or P300 peaks. As done in most oddball studies, one way is to select the conventional scalp locations at Fz, Cz and Pz and/or average them, but this does not sufficiently address the problem. Since our overarching goal is to elucidate the modality-specific cortical generators of MMN and P300, we employed a data-driven approach and identified the sensor-space electrodes with maximum contribution to the MMN and the P300 activities in each condition. Using the MATLAB functions *min* and *max* on the difference waveforms, we first computed the top 5 sensors contributing individually to the MMN and P300 peaks, respectively. The top 5 sensors showing maximum MMN activity (maximum negative voltage) were F1/z/2, FCz/2 in the *audio only* condition; P1/3, PO3/5/7 in the *visual only* condition; P1/z/2/4, PO3 in the *AV* condition; POz/3/4/5, Oz in the *cross-audio* condition and CP1/3/5, P1/3 in the *cross-visual* condition. Similarly, electrodes F1/z/2, FC1/z showed maximum P300 activity in the *audio only* condition; electrodes FC1/z, Cz, Cpz, Pz showed highest activity in the *visual only* condition; electrodes Fz, FC1/z, Cz, Pz in the *AV* condition; electrodes F1/z/2, FC1/z in the *cross-audio* condition and electrodes F1/z, FC1/z/2 showed maximum P300 activity in the *cross-visual* condition. To deal with the multiple comparisons problem of dealing with different set of electrodes for different conditions, we reduced the dimensionality across the sensor space using Principal Component Analysis (PCA). We employed the *pca* function from MATLAB on the subject-level difference waveforms of MMN and P300 for each condition with their respective top 5 sensors. PCA computes the eigenvectors of the covariance matrix (“principal axes”) and sorts them by their eigenvalues (amount of explained variance). The “loadings” obtained as outputs of the *pca* algorithm were extracted to create a spatial filter for each condition which was projected to the individual standard and oddball waveforms. A common spatial filter was used here to optimize the differences between the standard and deviant categories within a given condition. **Figure 2** illustrates the “scores” of the first three principal components as topoplots along with their corresponding eigenvalues (λ) in the principal component subspace. As evident, the first principal components in all conditions captured the maximum variance and alone explained more than 80% of the data. Therefore, to better characterize the amplitude and latencies of the prediction error markers across different modalities, we used the data corresponding to the first PC only.

**Figure 2.**
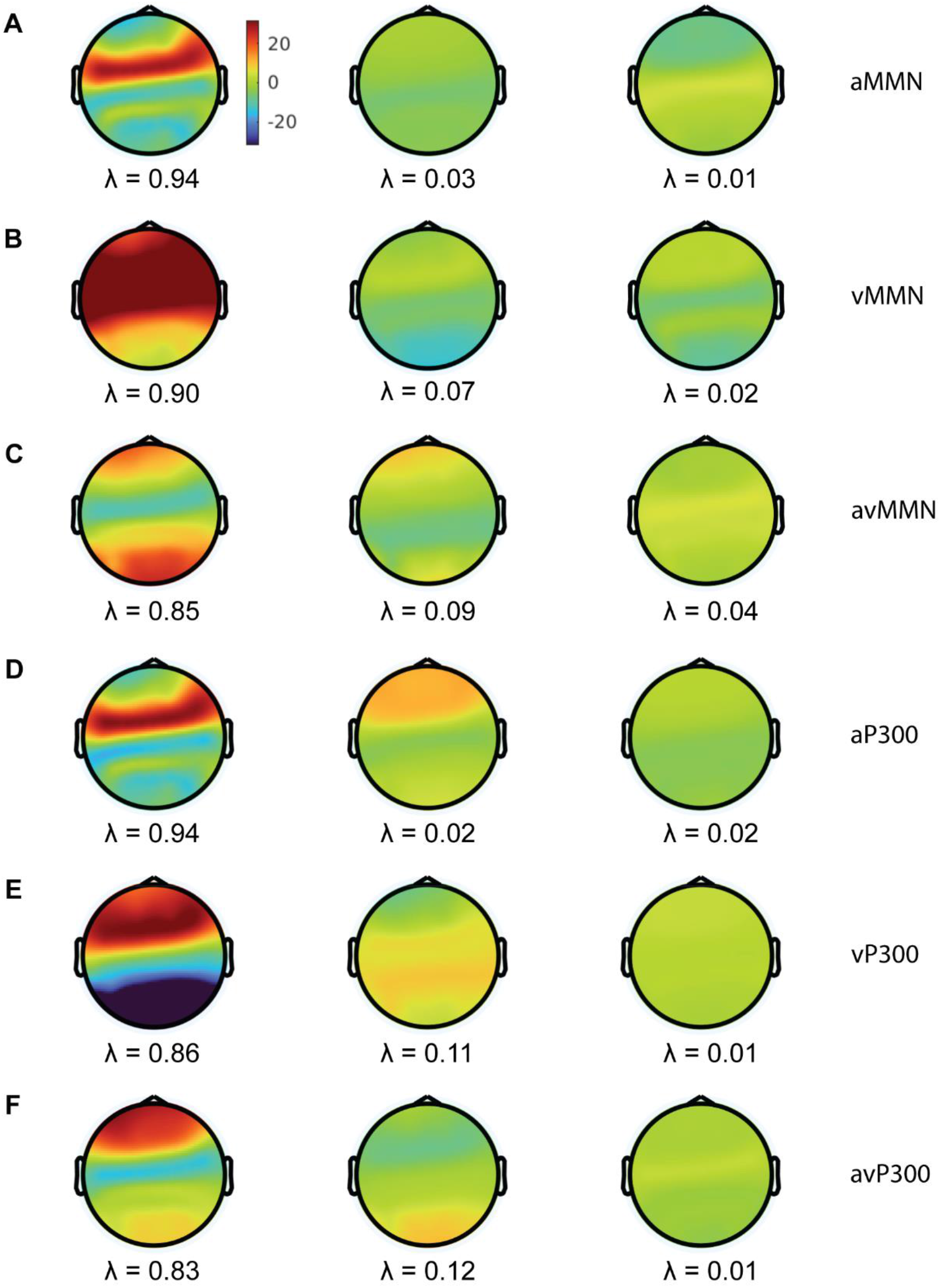
The topoplots correspond the first three principal components (left to right) of the spatial filters computed through PCA on the difference waveforms of the ERP **components (A) aMMN, (B) vMMN, (C) avMMN, (D) aP300, (E) vP300** and **(F) avP300** extracted using the standard and deviant waveforms from the top five sensors contributing to the MMN and P300 peaks in the audio, visual and audio-visual conditions. The λ values represent the variance explained by each principal component.

Note:- For a detailed statistical account on the amplitudes and latencies of MMN and P300, we mainly considered the *audio only, visual only* and *AV* conditions only due to the unavailability of ‘equivalent standard categories’ to subtract from the other two *cross-modal* deviant conditions. However, the two *cross-modal* conditions have been incorporated later in the study to investigate the robustness of the source distribution results.

### Source Localization

Source localization was performed to obtain the MMN and P300 generators in the brain across various modalities. The EEG data were re-referenced with an average reference prior to source-localization. 19 participants’ structural MRI data were re-sliced and segmented to identify the brain, skull and scalp tissues. 2 subjects’ MRI scans could not be obtained because of their incompatibility with the fMRI scanner. The origin (0,0,0) of all the T1 images was set to the anterior commissure. Participant-specific headmodel was computed using the OpenMEEG toolbox (Gramfort et al., 2010), using realistic conductivity values. The Polhemus data was imported to place the sensor locations on the head model of each participant. To obtain high accuracy of electrode positions, individual co-registration was employed by firstly visually marking the fiducials (nasion, left preauricular and right preauricular) in the MRI volume and finally matching the marked points with the fiducial locations as per the Polhemus data. Next, the sources were placed in the segmented part of the headmodel at a distance of 5 mm from each other and the leadfield matrix was computed, i.e., a transfer function between each source point and each electrode. Source localization of each individual was performed using their respective headmodel, leadfield and electrode positions. eLORETA, belonging to a class of current density measures to calculate the distribution of neuronal sources, was used to solve the inverse problem (Pascual-Marqui, 2007). eLORETA also generates the least amount of false positives hence beneficial for exploratory source analysis, e g., where prior hypotheses regarding approximate locations may not be available (Halder et al., 2019). Lambda of 0.1 was used as the regularization parameter for localization of P300 and MMN ERPs. After localization, each individual’s source intensities were interpolated to their respective MRI volume. Further, to calculate the grand average of the source values, the interpolated images were normalized to a common structural template. Finally, we subtracted the voxel intensities of oddball and standard categories and the voxels having intensities more than 95 percent of the maximum value were thresholded. This was done separately for each hemisphere.

### Data availability and codes

Codes used to analyze the data and reproduce the figures can be obtained from the bitbucket repository using the following link: https://bitbucket.org/cbdl/pe_erpandsourceanalysis/src/master/.

EEG data would be made available by the authors for public access upon acceptance of the manuscript. Copyright of the data and codes are held by the National Brain Research Centre, an autonomous institution of the Government of India.

## RESULTS

### Middle and late stages of prediction error processing conserved across unisensory and multisensory contexts

We inspected the subject-wise data (**Figure 3**), post projection of the common spatial filter on the standard and deviant waveforms. The difference waveforms (deviant – standard) revealed clear MMN (most negative peak between 120-250ms) and P300 (most positive peak between 250-500ms) responses (**Figure 3**) in the audio only (A), visual only (V) and audio-visual (AV) conditions. In each condition, to statistically verify the presence of the P300 peaks and the more negative MMNs in deviant trials as compared to the standards, the standard and deviant waveforms of each participant were averaged over the time-points within 120-250 ms for MMN and 250-500 ms for P300 which were subjected to a Student’s t-test. Even though the audio only (A) and audio-visual (AV) standards and deviants were found to be statistically different (p<0.001) for both the MMN and P300 latencies, the visual only (V) standard and deviant waveforms showed statistical difference (p<0.001) only for MMN. A prominent visual P300 can be seen in **Figure 3E (bottom-panel)** but since the P300 peak appears comparatively late and is preceded by a negative peak, the deviant waveform does not show any significant difference from the standard when the entire P300 temporal window (250-500 ms) was under consideration. This necessitated the use of modality-specific windows centered around the group-averaged peak (±50ms) for each condition. To maintain uniformity in the variances of the ERP peaks, equal window lengths of 100 ms were computed for both MMN and P300 and Student’s t-test was re-run on these newly computed windows. The 100 ms windows were sufficient to include the descending and ascending limbs of MMN peaks and the ascending and descending limbs of the P300 peaks. The t-test results showed that the amplitude of MMN in the group-averaged deviant trials was significantly lower than that of the standard trials in *audio only* (158-257ms : t(20)=-5.94, SD=4.88, p<0.001), *visual only* (109-208ms : t(20)=-5.90, SD=3.01, p<0.001) and *audio-visual* (102-201ms : t(20)=-6.52, SD=3.91, p<0.001) conditions (**Figure3A-C, bottom panels**). Similarly, now the P300 amplitudes were significantly higher for the deviant waveforms in *audio only* (281-380ms : t(20)=9.87, SD=5.59, p<0.001), *visual only* (353-452ms : t(20)=6.15, SD=4.13, p<0.001) and *audio-visual* (281-380ms : t(20)=6.28, SD=7.76, p<0.001) conditions (**Figure3D-F, bottom panels**).

**Figure 3.**
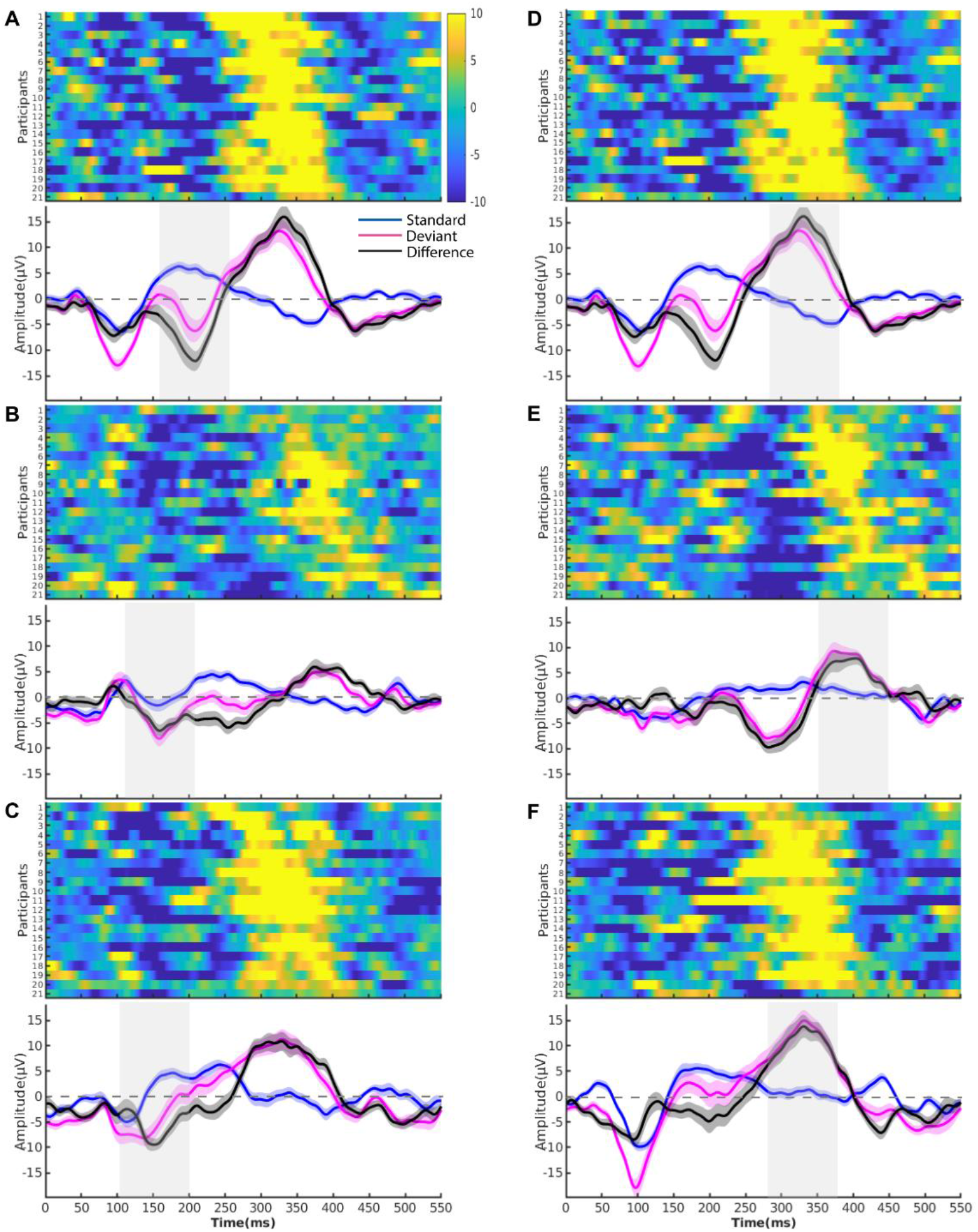
The heatmaps (top-panels) depict the time-course of difference waveforms of individual participants after projection of the common spatial filter (obtained through PCA) on the standard and deviant data, presented with the grand-averaged plots (bottom-panels) of the standard (blue), deviant (magenta) and difference (black) waveforms along with their shaded SEMs for **(A) aMMN, (B) vMMN, (C) avMMN, (D) aP300, (E) vP300**, and **(F) avP300**. The shaded grey windows in the bottom panels represent a regime of significant difference between the standard and deviant waveforms calculated using a Student’s t-test. Equal 100ms windows centred around the MMN (maximum negative activity) and P300 (maximum positive activity) peaks were considered to minimize the variance in the data across conditions.

### Multisensory context speeds the processing of prediction errors

The minimum and maximum amplitudes of each subject and their corresponding latencies were extracted for amplitude and latency characterization of MMN and P300, respectively from the difference waveforms (deviant-standard) across audio only (A), visual only (V) and audio-visual (AV) conditions. The *min* and *max* functions of MATLAB were used for this purpose. Subsequently, the group means and standard deviations of latencies and amplitudes were obtained for all conditions as listed in **Table 2** for P300 and in **Table 3** for MMN.

**Table 2.** Table shows the mean±SD of the peak P300 amplitudes and their corresponding latencies across all the subjects for audio only, visual only and audio-visual conditions.

| *Condition (P300)* | *Latency (ms)* | *Amplitude ($\mu V$)* |
| --- | --- | --- |
| *Audio only* | 336.57±25.85 | 20.42±9.47 |
| *Visual only* | 387.43±24.23 | 13.56±7.98 |
| *Audio-visual* | 332.57±26.87 | 18.23±9.66 |

**Table 3.** The table shows the mean±SD of the peak MMN amplitudes and their corresponding latencies across all the subjects for audio only, visual only and audio-visual conditions.

| *Condition (MMN)* | *Latency(ms)* | *Amplitude($\mu V$)* |
| --- | --- | --- |
| *Audio only* | 200.90±24.64 | 17.18±8.21 |
| *Visual only* | 161.43±31.33 | 10.67±3.63 |
| *Audio-visual* | 153±24.92 | 13.82±5.59 |

We systematically investigated the interaction of modalities with the latencies and amplitudes of MMN/P300 peaks, by conducting a repeated-measures analysis of variances (rmANOVAs) on the subject-level data. The results revealed a significant effect of modality on the latencies of both MMN (F(2,40)=23.421, p<0.001, η^2^=0.54) and P300 (F(2,40)=41.075, p<0.001, η^2^=0.67); and also on the amplitudes of MMN (F(2,40)=7.301, p=0.002, η^2^=0.27) and P300 (F(2,40)=8.104, p=0.001, η^2^=0.28). If rmANOVA is not used, a simple ANOVA would inflate Type I error (false-positives) in between-subject effects and Type II error (false-negatives) in within-subject effects. To learn more about various univariate measures and their impact on Type I and Type II errors, refer to Fields & Kuperberg, 2020. Another important point to keep in mind while using univariate rmANOVA is that the variances of the differences between all combinations of measurements should be equal. This is known as the sphericity (or circular) assumption which is strongly assumed for within-subject rmANOVA statistics. If the sphericity assumption is violated, within-subject rmANOVA statistics are meaningless. We used Mauchly’s test to verify that the assumption of sphericity was not violated (p>0.05) in any of our conditions. Furthermore, to employ a multiple comparisons correction that provides a good statistical power, post-hoc analyses were performed using Bonferroni test to compare the peak MMN and P300 latencies/amplitudes between different conditions. The peak amplitudes in the audio only condition were significantly higher than the peak amplitudes in the visual only condition for both MMN (p=0.011) and P300 (p=0.001). The peak MMN/P300 amplitudes of the AV condition however, did not show any significant difference from peak amplitudes of the audio only (p=0.229 for MMN, p=0.470 for P300) as well as the visual only (p=0.059 for MMN, p=0.104 for P300) conditions and were intermittent to the MMN/P300 peak amplitudes of both the conditions. The corresponding P300 latencies were shortest (and fastest) for the AV and audio only conditions (with no significant difference between the two conditions) and highest (and slowest) for the visual only condition (V>AV and V>A at p<0.001). The corresponding MMN latencies were shortest for the AV and the visual only conditions (AV<V at p=0.214) while the average MMN latency was largest in the audio only condition (A>V and A>AV at p<0.001).

### Spatial representations of unisensory and multisensory contexts in source space

The 100 ms windows of each modality were used to obtain the covariance matrices, separately for standard and oddball trials of MMN. The clusters of P300 sources obtained using 100 ms windows, however, were noisy (did not fall into any Brodmann area), because of which we recomputed the P300 sources using a longer time window to obtain a better estimate of variance in the data. Thus, the P300 peaks were identified between 250-500 ms post-stimulus onset where maximum peak values of every trial were extracted along with their corresponding latencies. Based on these latencies, we calculated the mean and standard deviation for each modality and defined the windows as [mean-SD : mean+SD]. The covariance matrices were now obtained from these windows and the same steps were repeated to obtain the P300 sources. Using co-registration of individual participant MR data with their EEG sensor locations, we generated the source maps underlying MMN and P300 activity using eLORETA (details in Methods). Parcels from the Brainnetome atlas were interpolated to the same common structural template as used for normalization of the individual sources. Only those parcels were chosen which included the sources or a part of them. As revealed in **Figure 5A**, the MMN sources were distributed throughout the brain and were different for different modalities (locations in **Table 4**).

**Table 4.**
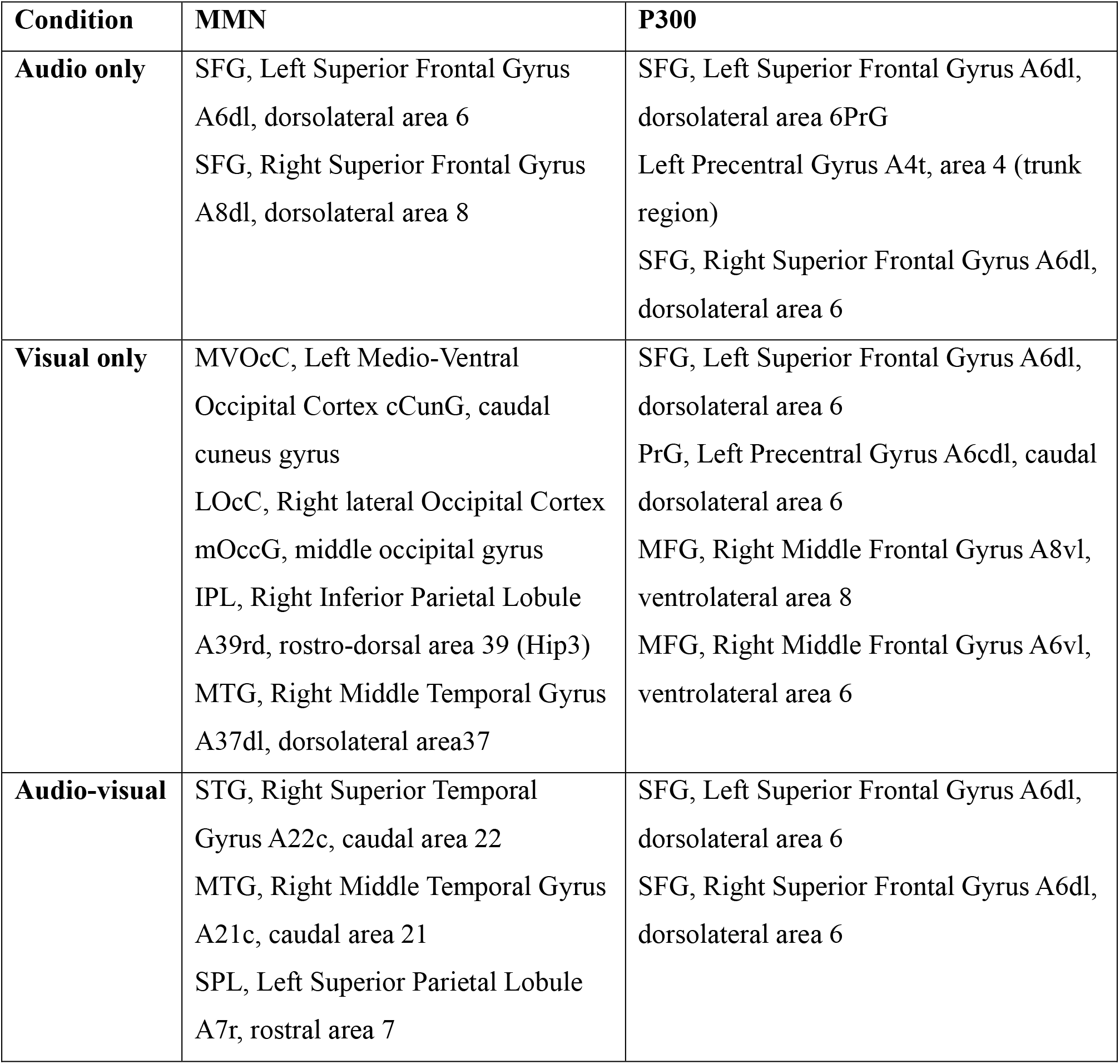
The table lists the brain areas underlying the peak MMN and P300 activations across the audio only, visual only and audio-visual conditions.

| **Condition** | **MMN** | **P300** |
| --- | --- | --- |
| **Audio only** | SFG, Left Superior Frontal Gyrus A6dl, dorsolateral area 6<br>SFG, Right Superior Frontal Gyrus A8dl, dorsolateral area 8 | SFG, Left Superior Frontal Gyrus A6dl, dorsolateral area 6PrG<br>Left Precentral Gyrus A4t, area 4 (trunk region)<br>SFG, Right Superior Frontal Gyrus A6dl, dorsolateral area 6 |
| **Visual only** | MVOcC, Left Medio-Ventral Occipital Cortex cCunG, caudal cuneus gyrus<br>LOcC, Right lateral Occipital Cortex mOccG, middle occipital gyrus<br>IPL, Right Inferior Parietal Lobule A39rd, rostro-dorsal area 39 (Hip3)<br>MTG, Right Middle Temporal Gyrus A37dl, dorsolateral area37 | SFG, Left Superior Frontal Gyrus A6dl, dorsolateral area 6<br>PrG, Left Precentral Gyrus A6cdl, caudal dorsolateral area 6<br>MFG, Right Middle Frontal Gyrus A8vl, ventrolateral area 8<br>MFG, Right Middle Frontal Gyrus A6vl, ventrolateral area 6 |
| **Audio-visual** | STG, Right Superior Temporal Gyrus A22c, caudal area 22<br>MTG, Right Middle Temporal Gyrus A21c, caudal area 21<br>SPL, Left Superior Parietal Lobule A7r, rostral area 7 | SFG, Left Superior Frontal Gyrus A6dl, dorsolateral area 6<br>SFG, Right Superior Frontal Gyrus A6dl, dorsolateral area 6 |

The dorsolateral areas of the left and right superior frontal gyri were found to underlie the MMN response in the *audio only* condition; the caudal cuneus gyrus in medio-ventral occipital cortex on the left, the middle occipital gyrus in the right lateral occipital cortex, the rostro-dorsal area of the right inferior parietal lobule and the dorsolateral area of the right middle temporal gyrus elicited MMN in the *visual only* condition; and the *audio-visual* condition yielded sources that were located in the rostral areas of left superior parietal lobule and the caudal areas of multisensory regions STG (superior-temporal gyrus) and MTG (medial-temporal gyrus) on the right. Interestingly, source analysis of P300 revealed many overlapping fronto-central areas for all the three sensory modality conditions (as illustrated in **Figure 5B**). P300 was elicited at the dorso-lateral areas of left superior frontal gyri for *audio only, visual only* and *audio-visual* conditions; at right superior frontal gyri for the *audio only* and *audio-visual* conditions; at the trunk and caudal dorso-lateral areas in left precentral gyri for the *audio only* and *visual only* conditions, respectively; and additionally, at the ventro-lateral area of right middle frontal gyrus for the *visual only* condition (details in **Table 4**).

## DISCUSSION

To detect neural activity with high temporal precision and distinguish between different neuro-cognitive processes on the basis of differences in scalp distribution is key to the ability of EEG to reveal the temporal stages of prediction error processing. We employed two-stimuli oddball tasks using *Audio alone, Visual alone* and *synchronous Audio-Visual* conditions to evoke MMN and P300 responses in the brain, considered as ERP level markers of prediction error processing in the literature (Aitchison & Lengyel, 2017). It is already known that visual modality speeds up auditory processing for both speech and non-speech stimuli (Begau et al., 2021; Diaconescu et al., 2011; Leone & Mccourt, 2015; van Wassenhove et al., 2005), previously demonstrated mostly using speeded N100 latencies in multisensory (audio+visual) contexts. Our study is an advancement to these previous findings and our first result elucidates how the simultaneous occurrence of audio and visual oddballs, shorten the latencies of both MMN and P300 ERPs, indicating that the processing of prediction errors gets facilitated under multisensory contexts at “any stage” of information processing hierarchy in the brain (early N100, middle MMN or late P300). This information is crucial and can serve as a powerful tool to formulate “internal models” trying to understand prediction error processing at different stages/hierarchies in the brain (Wacongne et al., 2012). No single study before ours has demonstrated this so explicitly involving both the prediction error markers using a common oddball paradigm in healthy humans. The second major outcome of our study in the context of prediction error processing is the modality-specific and modality-agnostic nature of the MMN and the P300 sources, respectively. This is the first study as per our knowledge to use accurate source localization techniques to report all the underlying areas of MMN and P300 in audio, visual and AV modalities. We found a reorganization of brain regions along the temporal hierarchy of prediction error processing (MMN followed by P300) as the mismatch information flows up the spatial hierarchy of the brain (sensory to higher order regions). This was revealed by an initial “modality-sensitive” stage indexed by MMN where different modalities had different cortical generators, which transitioned to a “modality-independent” stage indexed by P300 where the different modalities had common cortical sources (**Figure 5, 6**). Though such a transition may appear unsurprising in terms of neuro-cognitive processing, we questioned at what stage in the information processing hierarchy is the modality specificity to the sensory stream lost? We found that the processing of prediction error becomes amodal at the P300 level itself and is not something that is attained at further later stages during the reorienting negativity (RON). The RON in the brain appears as a negative deflection at 400-600ms in response to the reorientation of attention to the predictive/standard stimuli after it has been switched (indexed by P300) towards the preceding deviant stimuli (Correa-Jaraba et al., 2016; Ungan et al., 2019). Even though in our analysis we could not cover this ERP since it did not come out as a prominent peak in the subtraction waveforms (**Figure 1**), we argue that RON too would, being an even later component, have common brain generators across modalities, probably more frontal, as also reported in recent studies (Correa-Jaraba et al., 2016; Justo-Guillén et al., 2019; Ungan et al., 2019). One may also note here that in our study, we have specifically focussed on only the two ERPs of interest-MMN and P300, as they have been widely reported as the markers of prediction errors across various modalities in extant literature (Banellis et al., 2020; Calcus et al., 2015; Chennu et al., 2013; Stefanics et al., 2014). Other prominent peaks in response to auditory stimuli such as N100 (as seen in **Figure 1A, 1C**), are out of the scope and interest of this study.

We compared the MMN and P300 latencies during the processing of prediction errors across audio only (A), visual only (V) and audio-visual (AV) conditions. The MMN latencies in our oddball paradigm followed the order: *Audio-visual<Visual only<Audio only* (**Table 3**). According to our results, the aMMN appears much later than the vMMN as also reported earlier in a few studies (Berti & Schröger, 2001; Sittiprapaporn, 2012). The MMN latencies are reported to be higher for smaller deviances (Horváth et al., 2008), and since the subjective magnitude of auditory deviance was small (low pitch vs high pitch) as compared to the magnitude of visual deviance (blue squares vs red triangles), the MMN onset latency for auditory deviants was evoked relatively later. We speculate that the delayed aMMN might also be because of the time taken for the sensory mismatch information to pass from one hierarchal cortical stage to the next, up to the frontal gyrus (**Table 4**). vMMN sources were in the occipital cortex whereas aMMN was elicited at a comparatively higher hierarchy in the superior frontal gyrus (as also reported previously by Näätänen et al., 2007; Winkler et al., 1998 in the sensor space). Because of the overlapping cortical generators of aMMN and aP300, bottom-up information flow time for further auditory prediction error processing post MMN elicitation was reduced. Consequently, the audio only condition produced the slowest MMN but the fastest P300 peaks (**Figure 4D**). The P300 latencies followed the order: *Audio-visual<Audio only<Visual only* (**Table 2, Figure 4D**). Superior-frontal auditory MMN might be indicative of a stronger call for attention towards change in a single stimulus feature i.e., frequency change in tone, that perhaps would require an active comparison of the mismatch with the standard to be identified as a deviant, as opposed to changes in basic visual features like color/shape in the visual oddball condition. This is supported by evidence from literature that suggests object-based irregularities are automatically detected by the visual system (Stefanics et al., 2014). MMN latency speeds up with increasing deviation from the standard (Näätänen et al., 2005) as the mismatch between the deviant and the memory representation of the standard can be detected faster. In our paradigm, the deviation in the visual oddball (change in shape and color) was high (or rather obvious), resulting in a faster processing speed of vMMN, and was resolved at the level of the secondary sensory cortex itself. However, visual mismatch information had to travel from the occipital cortex higher up to the superior-frontal gyrus for the next level of mismatch processing at P300 latency, resulting in slowest vP300. We found this result quite intriguing because the MMN and P300 latencies in the temporal hierarchy organized themselves based on the arrangement of brain areas along the spatial hierarchy of the predictive brain supporting previous studies stating that MMN may be influenced by recurrent feedback activation from higher order areas (Garrido et al., 2007, 2009). More importantly, fastest elicitation of avMMN (**Figure 4C**) suggests that the faster visual component in the AV oddball temporally facilitated its slower auditory component which speeded up the process of change detection in the audio-visual condition. Evidently, this facilitation for audio-visual deviants was propagated further up the deviant processing ladder resulting in fastest avP300 localized at the superior-frontal gyrus. Both avP300 and aP300 were significantly faster than vP300, suggesting a temporal facilitation of audio-visual deviants at the level of P300 latency. Taken together, our results clearly reveal that the presence of a multisensory context, shifts the oddball processing speed towards the faster unisensory modality indicating a multisensory benefit at “all stages” of processing prediction errors.

**Figure 4.**
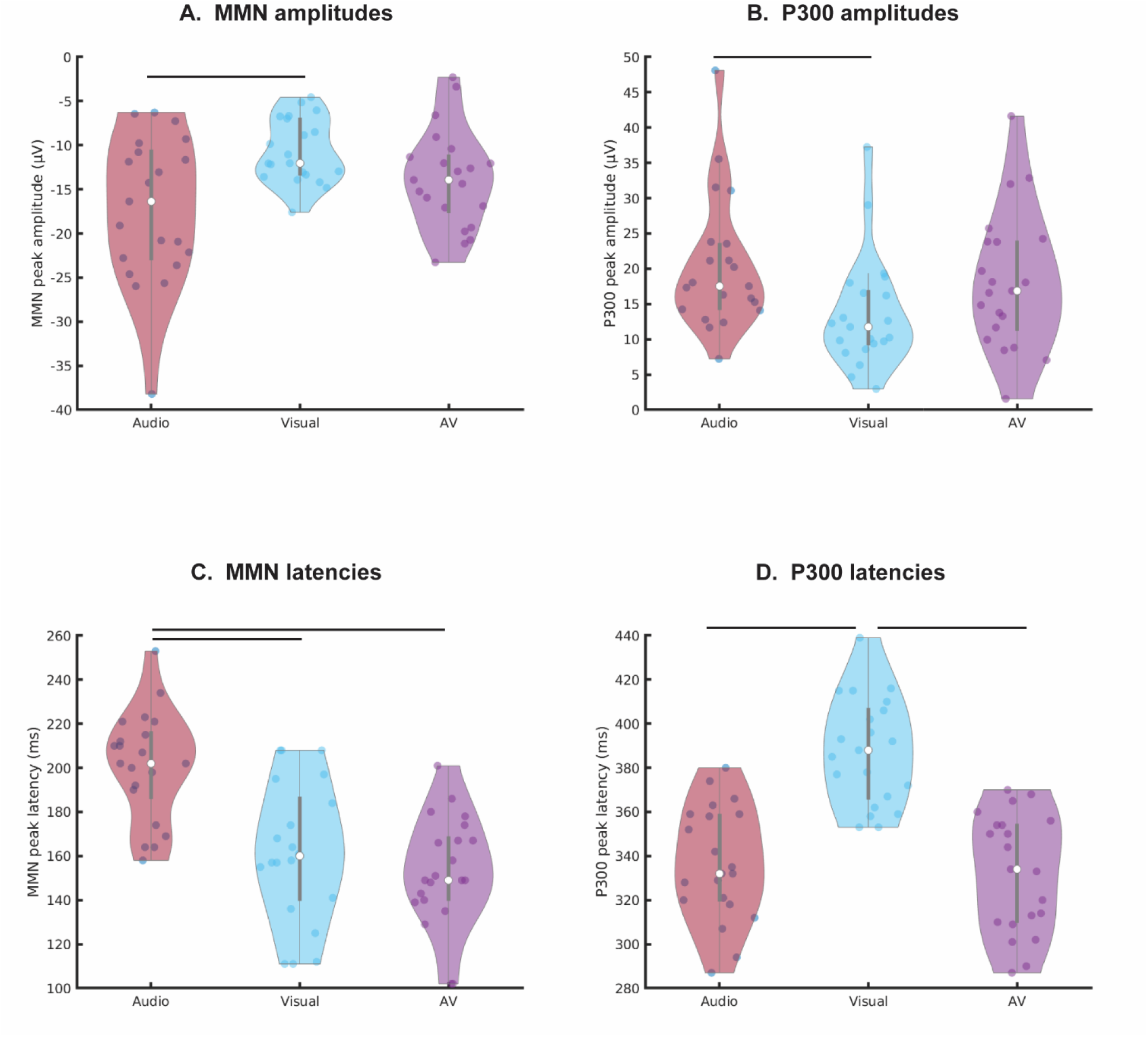
The violin plots represent the **(A) MMN peak amplitudes** (maximum magnitude of the difference waveforms on the negative y axis) of each participant and the corresponding **(C) MMN peak latencies**. Similarly, the **(B) P300 peak amplitudes** and the corresponding **(D) P300 peak latencies** are plotted for each participant. The colored dots represent each participant’s peak amplitude and peak latency values in µV and ms, respectively. The white dot at the center of the gray box in each violin represents the median of the data and the gray box itself represents the inter-quartile range. The horizontal lines represent significant difference (p<0.05) between any two conditions obtained through Bonferroni corrected post-hoc analysis.

**Figure 5.**
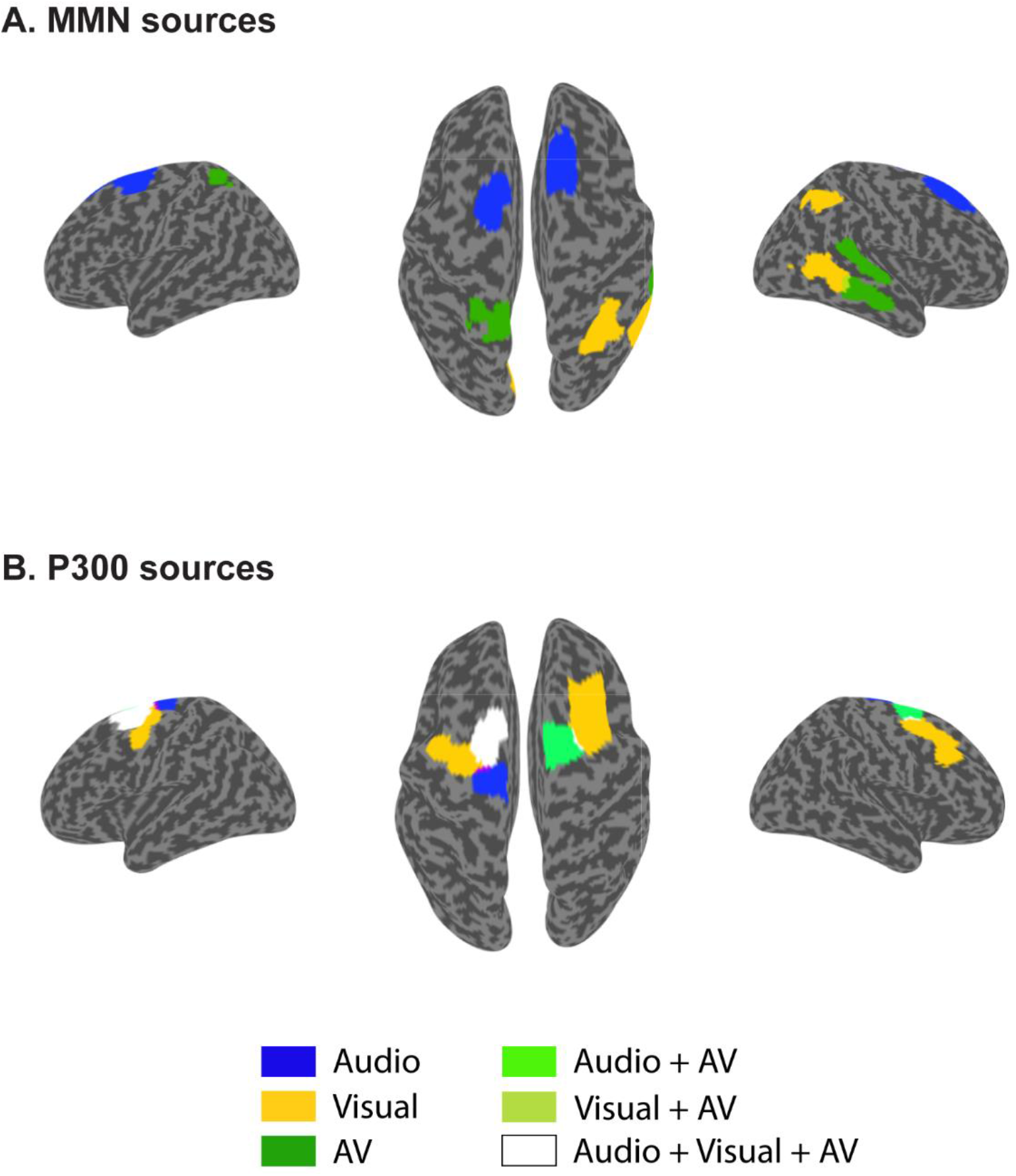
eLORETA source localization results using time-locked analysis (at a threshold of 95%) representing the underlying **(A) MMN** and **(B) P300 sources** for Audio, Visual and Audio-Visual modalities.

One interesting possibility underlying the speeded appearance of the audio-visual prediction markers can be an early coupling in the information processing hierarchy between the constituent unisensory audio and unisensory visual deviant stimuli during an early sensory processing stage (e.g., at N100). An alternate account could be that the unisensory systems operate largely independently to extract feature-level information, before multisensory convergence and integration marked by audio-visual MMN at the superior-temporal gyrus and middle temporal gyrus, the higher order association areas (Beauchamp, 2005; Callan et al., 2004). Earlier studies on multisensory MMN related to somatosensory and auditory modalities argued that if auditory and somatosensory information interacted to influence perception during the sensory-perceptual stages of processing, the MMN in response to combined auditory and somatosensory deviants would display multisensory integrative properties (Butler et al., 2011, Butler 2012). Alternatively, if there was no intersensory coupling at the sensory-perceptual level, the MMN for combined somatosensory and auditory deviants should be reflecting a linear summation of the individual auditory and somatosensory MMNs (Molholm et al., 2002). In our results however, we saw a clear non-linearity in the multisensory response where the amplitudes of the avMMN and avP300 were smaller than the sum of aMMN and vMMN as well as that of aP300 and vP300, respectively (**Table 2, 3**). The amplitudes of the aMMN and vMMN (as well as aP300 and vP300) differ significantly from each other (aMMN>vMMN and aP300>vP300) and the amplitude of avMMN (and avP300) is intermediary to them, reflecting the sub-additive nature of the multisensory effect at all stages of prediction error processing hierarchy. This further suggests that the aMMN and aP300 responses were minimized when paired with the concurrent visual stimulation which can be attributed to the bimodal nature of our audio-visual stimuli as the presence of two stimuli gives the brain more flexibility to choose deviant features to process the AV oddballs efficiently and hence, conserve its neural resources during change detection in accordance to the free energy principle (Friston, 2010).

An important aspect of our present study is that the hierarchical stages of prediction error processing, can be explained from multiple underlying internal models of error propagation (Aitchison & Lengyel, 2017). Since, the synchronous presentation of the audio and visual deviants in our paradigm may not form any unitary multisensory construct like in case of speech stimuli (or even non-speech ventriloquist like stimuli as demonstrated by Stekelenburg et al., 2004) and thus identifying the AV deviants from a stream of AV standards could easily be achieved by only selectively attending to the most salient feature of the audio-visual oddballs, for instance, change of color from blue to red instead of any underlying free energy minimization process. To minimize refractoriness in the sensory cortical feature specific neurons (Jacobsen & Schröger, 2001), there may also be a switch between which features the brain attends to during the course of the oddball trials. It may be noted here that the observed multisensory benefit in our paradigm in case of avMMN and avP300 can readily be attributed to a “multisensory interaction” in which the unisensory auditory and unisensory visual streams interact with each other due to temporal synchrony. This however may not be equivalent to “multisensory integration” where the multisensory stimuli forms a new perceptual meaning. There also exists evidence (Tsilionis & Vatakis, 2016) which suggests that multisensory integration and interaction (referred to as perception synchrony) involve separate mechanisms that are not tightly coupled because multisensory interaction may be immune to semantic relations relying on low-level characteristics, while multisensory integration, although influenced by timing, is critically dependent on semantics. In the future, to better understand the nature and extent of audio-visual interactions, it will be essential to design advanced paradigms that form meaningful stimuli in both unisensory and multisensory constructs. It may also be noted that the two auditory stimuli presented in the present paradigm, were pure tones with musical relation in terms of pitch (deviance was one octave higher, **Table 1**). This may indicate that the MMN and the P300 obtained for *audio only* condition represent the neural signatures of auditory pitch mismatch processing.

Source localization results for MMN showed heterogenous activations in the brain across different sensory modalities (as shown in **Figure 6A**). For the *visual only* modality, we saw activations in the left medio-ventral occipital cortex, the right lateral occipital cortex, the right inferior parietal lobule and the right middle temporal gyrus; for *audio only* condition, there were activations in the left and right superior frontal gyri; for the *audio-visual* modality, activations were seen in the left superior parietal lobule and the right superior-temporal and medial-temporal gyri; for the *cross-visual* modality, we found activations in the left and the right superior parietal lobules; and that for *cross-audio* modality, in the left and the right lateral occipital cortex (**Figure 6A**). From these results, we concur that mostly the secondary sensory areas along with other cortical areas distributed throughout the brain are employed during the process of early prediction mismatch, based on the modality. An interesting point to be noted here is that even though the deviant stimuli were exactly the same in the *Audio only* and *Cross-Audio* conditions (and similarly in the *Visual only* and *Cross-Visual* conditions), their cortical generators were different only because their corresponding standard stimuli were different suggesting that the MMN sources are not only sensitive to the modality of the deviant stimuli but also to the modality of the standards.

**Figure 6.**
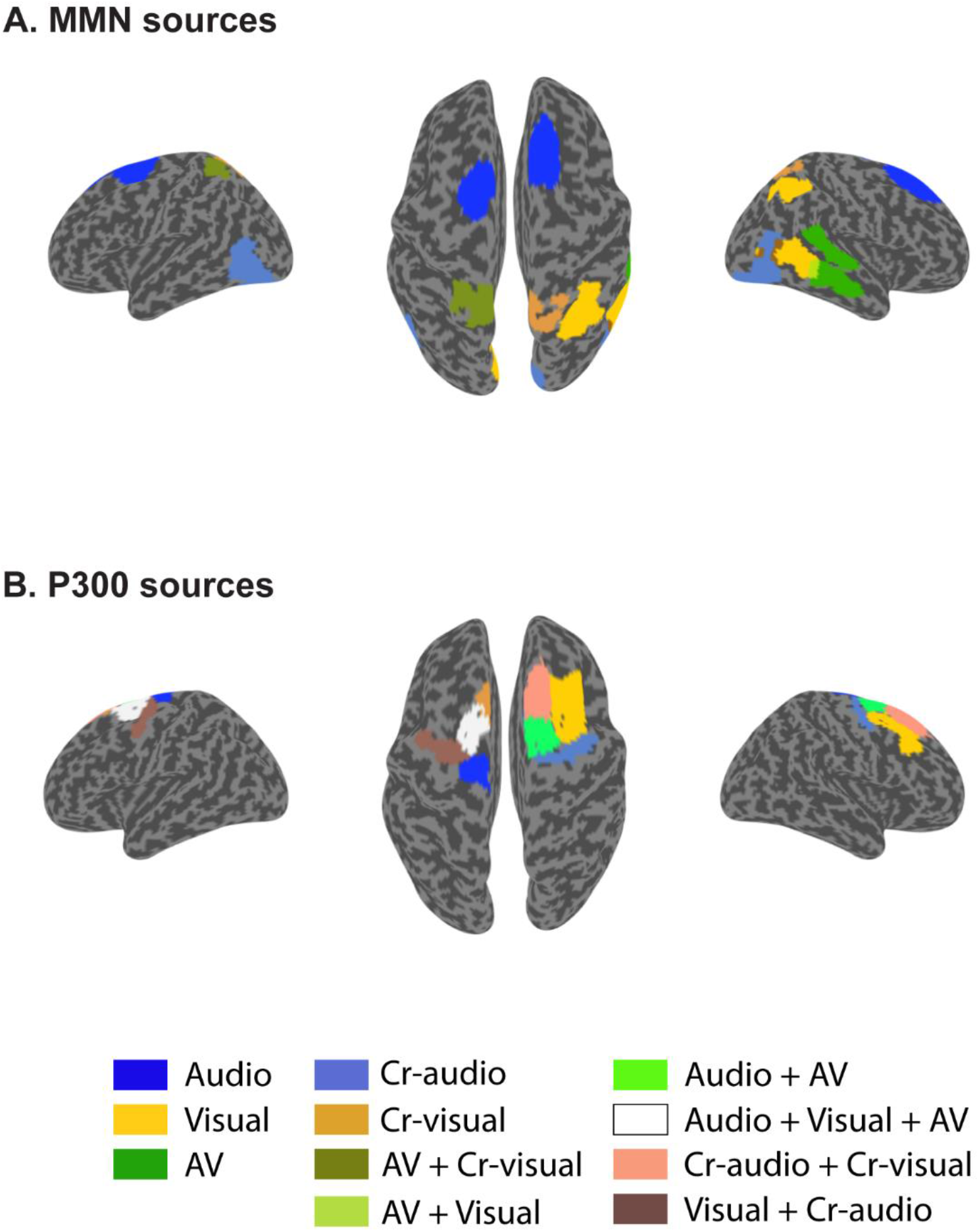
eLORETA source localization results using time locked analysis (at threshold level 95%) representing the underlying **(a) MMN** and **(b) P300 sources** for Audio, Visual, Audio-Visual, Cross-Visual and Cross-Audio modalities.

In traditional literature, MMN is widely considered as a “perceptual” prediction error signal carrying important novel information which initiates a call for further processing of prediction mismatches (Escera et al., 2000; Mäntysalo & Näätänen, 1987; Schröger, 1997). Such novel information should necessarily feed into the higher cognitive processing areas for further evaluation, as is reflected by the occurrence of the late event-related potential, P300 (Bledowski, Prvulovic, Hoechstetter, et al., 2004; David & Linden, 2005). Our source results revealed overlapping source activations for P300 in all the five modality conditions, localized around the fronto-central regions for *audio only, visual only* and *audio-visual* conditions (as shown in **Table 4**) and also at the right superior frontal gyrus, the left and the right precentral gyri for *cross-audio* condition and, at the left and the right superior frontal gyri for *cross-visual* condition (as shown in **Figure 6B**). Common brain generators for P300 across modalities suggest that this neural marker might particularly be responsible for domain-general higher evaluative processes like keeping a count of the number of oddballs as in our paradigm, after a change detection signal has been relayed by MMN generators of corresponding modalities in the brain.

Many recent and past studies have argued that the brain is not a passive input-output device but acts as a predictive system capable of anticipating the future by making predictions about sensory inputs and minimizing prediction errors (Bubic et al., 2010; A. Clark, 2013; Ficco et al., 2021; Friston, 2010b). Thus, there arises a need to further understand the neural mechanisms by which the brain processes prediction errors in complex scenarios and multiple modality combinations. A detailed neuronal model of the auditory, visual and audio-visual cortices, based on the underlying processes of predictive mismatch that account for the critical features of MMN, P300 and RON could better explain the process of temporal facilitation and supra-additivity in the audio-visual modality (Pattamadilok and Sato, 2022; Molholm et al., 2002). Our empirical investigations in the context of hierarchical processing of prediction errors would have implications beyond the theoretical domain as well. Based on the models of MMN and P300 responses from patients with disorders of consciousness like in vegetative and minimally conscious states (Boly et al., 2011; Daltrozzo et al., 2007), attention (Polich, 2007b; Szuromi et al., 2011), and schizophrenia (Blackwood, 2000; Erickson et al., 2016), researchers can isolate the deficits in predictive information flow that might underlie these states of profound cognitive and neurological dysfunction. Such foundational advances can be of extreme value to clinical neuroscience researchers. From a neural networks’ perspective, two well-known attentional networks, the dorsal attention network (DAN) and the ventral attention network (VAN) have been largely reported in oddball studies (Bledowski, Prvulovic, Goebel, et al., 2004; V. P. Clark et al., 2000; Stevens et al., 2000). The VAN in particular has been exclusively involved in the detection of deviant stimuli (Kim, 2014; Palaniyappan & Liddle, 2012) and is activated at both MMN and P300 stages (Justen & Herbert, 2018). Further connectivity analysis between the regions underlying the VAN during MMN and P300 latencies across various modalities can further our understanding of the function of this important attentional network beyond recent evidence in the frequency domain (Ghosh et al., 2021). Although a few studies have also attempted to draw a relationship between pre-stimulus brain states and P300 (Karch et al., 2016; Reinhart et al., 2011), a detailed source connectivity analysis will be informative about the mechanisms of P300 and MMN generation, thus providing an important direction to future research.

## Acknowledgments

We wish to acknowledge the support of NBRC core funds and the computational support from Computational Facility, NBRC. This project is funded by the NBRC Flagship program BT/ MEDIII/ NBRC/ Flagship/ Program/ 2019: Comparative mapping of common mental disorders (CMD) over lifespan.

## List of abbreviations

aMMN: auditory mismatch negativity
avMMN: audio-visual mismatch negativity
DAN: Dorsal Attention Network
EEG: Electroencephalography
eLORETA: Exact low resolution brain electromagnetic tomography
ERP: Event Related Potential
ICA: Independent Component Analysis
MRI: Magnetic Resonance Imaging
MTG: Medial Temporal Gyrus
PCA: Principal Component Analysis
rmANOVA: repeated measures Analysis Of Variance
RON: Reorienting negativity
STG: Superior Temporal Gyrus
VAN: Ventral Attention Network
vMMN: visual mismatch negativity

## AUTHOR CONTRIBUTIONS

PG, ST, and AB conceived the study; PG and ST designed the stimulus; PG collected the data and wrote the initial draft; PG and ST analyzed the data; PG, ST and AB revised the draft; AB supervised the work.

